# Pupil responses to pitch deviants reflect predictability of melodic sequences

**DOI:** 10.1101/693382

**Authors:** Bianco Roberta, Ptasczynski Lena Esther, Omigie Diana

## Abstract

Humans automatically detect events that, in deviating from their expectations, may signal prediction failure and a need to reorient behaviour. The pupil dilation response (PDR) to violations has been associated with subcortical signals of arousal and prediction resetting. However, it is unclear how the context in which a deviant occurs affects the size of the PDR. Using ecological musical stimuli that we characterised using a computational model, we showed that the PDR to pitch deviants is sensitive to contextual uncertainty (quantified as entropy), whereby the PDR was greater in low than high entropy contexts. The PDR was also positively correlated with unexpectedness of notes. No effects of music expertise were found, suggesting a ceiling effect due to enculturation. These results show that the same sudden environmental change can lead to differing arousal levels depending on contextual factors, providing evidence for a sensitivity of the PDR to long-term context.

## Introduction

The experience of surprise is very common in the sensory realm and often triggers automatic changes in arousal and attentional states that are fundamental to adaptive behaviours. Music is a ubiquitous and ecological example of a situation where changes in listeners’ arousal and attention are intentionally manipulated. Composers may, for example, modulate the predictability of musical passages in order to achieve differing levels of tension in a listener. A great deal of empirical work has shown that surprising sounds are recognised by listeners in an effortless and automatic fashion (Pearce, 2018). This process is thought to be supported by a mismatch between the current unexpected input and the implicit expectations made possible by schematic and dynamic knowledge of stimulus structure (Huron, 2006; Krumhansl, 2015; Tillmann, Bharucha, & Bigand, 2000; Vuust & Witek, 2014). However, there is still rather little research examining mismatch responses under different degrees of uncertainty during passive listening.

Evidence of listeners experiencing events in music as unexpected comes from studies investigating behavioural (Marmel, Tillmann, & Delbé, 2010; Omigie, Pearce, & Stewart, 2012; Tillmann & Lebrun-Guillaud, 2006) and brain responses to less regular musical events (Bianco, Novembre, Keller, Kim, et al., 2016; Carrus, Pearce, & Bhattacharya, 2013; Koelsch, 2016; Koelsch, Gunter, et al., 2002; Maess, Koelsch, Gunter, & Friederici, 2001; Miranda & Ullman, 2007; Omigie, Pearce, Williamson, & Stewart, 2013; Pearce, Ruiz, Kapasi, Wiggins, & Bhattacharya, 2010). With regard to the former, priming paradigms have shown that a context allows perceivers to generate implicit expectations for future events, leading to facilitated processing (i.e., priming) of expected events. With regard to the latter, violation paradigms have shown increased brain responses to deviant events (out of key notes, or harmonically incongruent chords) within structured contexts as well as events which are musically plausible but more improbable in the given context. For example, Omigie et al. (2013) tested brain responses to melodies whose notes were characterised in terms of their predictability by a model of auditory expectations (Pearce, 2005). They showed that surprising events (more improbable notes) within melodies elicited a mismatch response – often termed the mismatch negativity, MMN (Garrido, Kilner, Stephan, & Friston, 2009; Näätänen, Paavilainen, Rinne, & Alho, 2007). This response decreased in amplitude for progressively more predictable events, as estimated by a computational model of melodic expectation. A similar parametric sensitivity to note unexpectedness has since also been observed in subcortical regions like the anterior cingulate and insula (Omigie et al., 2019). Moreover, sensitivity to music structure violation seems to emerge in all members of the general population that have had sufficient exposure to a given musical system (Bigand & Poulin-Charronnat, 2006; Pearce, 2018; Rohrmeier, Rebuschat, & Cross, 2011), and this sensitivity is modulated by pre-existing schematic knowledge of music, as reflected in acquired levels of musical expertise (Fujioka, Trainor, Ross, Kakigi, & Pantev, 2004; Koelsch, Schmidt, & Kansok, 2002a; Tervaniemi, 2009; Vuust, Brattico, Seppänen, Näätänen, & Tervaniemi, 2012).

According to theoretical and empirical work framing perception in the context of predictive processing, the experience of surprise may be modulated by the predictability of a stimulus structure as it unfolds (Clark, 2013; Dean & Pearce, 2016; Friston, 2005; Ross & Hansen, 2016). Random or high entropic stimuli hinder the possibility of making accurate predictions about possible upcoming events, whilst stimuli characterized by familiarity or regular statistics will enable relatively precise predictions by permitting the assignment of high probability to a few possible continuations. Perceptually, it has been shown that listeners indeed experience high-entropic musical contexts with greater uncertainty compared to low entropy ones (Hansen & Pearce, 2014). Moreover, previous work has shown that neurophysiological signatures associated with auditory surprise display larger responses to a given deviant event when it is embedded in a low rather than high entropic context (Dean & Pearce, 2016; Garrido, Sahani, & Dolan, 2013; Hsu, Le Bars, Hamalainen, & Waszak, 2015; Ricardo Quiroga-Martinez, 2018; Rubin, Ulanovsky, Nelken, & Tishby, 2016; Southwell & Chait, 2018). Therefore, research suggests that to understand whether and how surprising events modulate arousal and re-orient behaviours, the statistics of the proximal context must be considered.

A vast literature has used pupil dilation response (PDR) as a general marker of arousal, selective attention, and surprise (Aston-Jones & Cohen, 2005; Sara, 2009). Pupil dilation is associated with the locus coeruleus-norepinephrine (LCN) system (Laeng, Sirois, & Gredeback, 2012; Widmann, Schröger, & Wetzel, 2018), the activation of which results in wide spread norepinephrine release in the brain. Increase of the PDR has been extensively reported in response to violation of expectations or surprising/salient events in distracted listeners (Damsma & van Rijn, 2017; Liao, Yoneya, Kidani, Kashino, & Furukawa, 2016; Wetzel, Buttelmann, Schieler, & Widmann, 2016; Zhao et al., 2019), and when deviants are presented below participants’ perceptual threshold (Fink, Hurley, Geng, & Janata, 2018). Furthermore, a relationship between PDR and continuous ratings of surprisal in music (measured via a continuous slider) has been reported during passive listening (Liao, Yoneya, Kashino, & Furukawa, 2018). Other work has specifically associated the PDR to violations of statistical regularities in the sensory input even when these violations are behaviourally irrelevant (Alamia, VanRullen, Pasqualotto, Mouraux, & Zenon, 2019; Zhao et al., 2019). These results provide supporting evidence for a role of norepinephrine in the tracking of abrupt deviations from the current predictive model of the world, and as a signal of prediction resetting that enables the discovery of new information (Dayan & Yu, 2006).

Music has the ability to play with our expectations, hence manipulating our arousal and emotions in an automatic fashion (Laeng, Eidet, Sulutvedt, & Panksepp, 2016; Meyer, 2001; Zatorre & Salimpoor, 2013). Abrupt changes in register, texture and tonality are all examples of instances where the listener may have to reset or potentially abandon current models about the unfolding music. Such changes may however appear less surprising if embedded in high entropy contexts. Here, we use music as an ecological setting with which to study how the PDR to deviant musical events is modulated by structure of the stimulus context, specifically by its entropy. The growing field of computational musicology means that information in melodies can be statistically estimated. One particular model of auditory expectations – the Information Dynamics of Music, IDyOM (Pearce, 2005) – has been shown to model listeners’ experience of surprise and uncertainty. This unsupervised Markov model learns and estimates the conditional probability of each subsequent note in a melody based on a corpus on which it has been trained (long-term sub-model; extra-opus-learning) and the given melody as it unfolds (short-term sub-model; intra-opus learning). The model outputs information content and entropy values, which, respectively, reflect the experienced unexpectedness of a certain note after its onset and the experienced uncertainty in precisely predicting a subsequent note based on the preceding pitch probabilities.

We created novel melodies (Figure 1) that adhered to the principles guiding Western tonal melodic structure. We then created shuffled versions of these melodies to create stimuli that were higher in entropy albeit matched for pitch range, content and tonal space. The information theoretic properties of all stimuli were estimated using IDyOM (Pearce, 2005). Listeners were presented with these melodies either in their standard form or with a pitch deviant whilst PDR was measured. Participants were not informed about the presence of the pitch deviants, but asked to rate the unexpectedness of last note of each melody. We expected a larger PDR to deviant notes that are embedded in low rather than high entropy contexts, and that are higher in their unexpectedness – information content value – as estimated by the computational model. Also, we expected entropy of the melodies to predict subjective ratings of stimulus unexpectedness (Hansen & Pearce, 2014). Finally, based on evidence of greater brain response to musical violations in musicians than non-musicians (Vuust, Brattico, Seppänen, Näätänen, & Tervaniemi, 2012b), presumably reflecting expertise-related enhancement in the accuracy of predictive models, differences due to musical expertise (Müllensiefen et al., 2014) were also investigated.

**Fig. 1.**
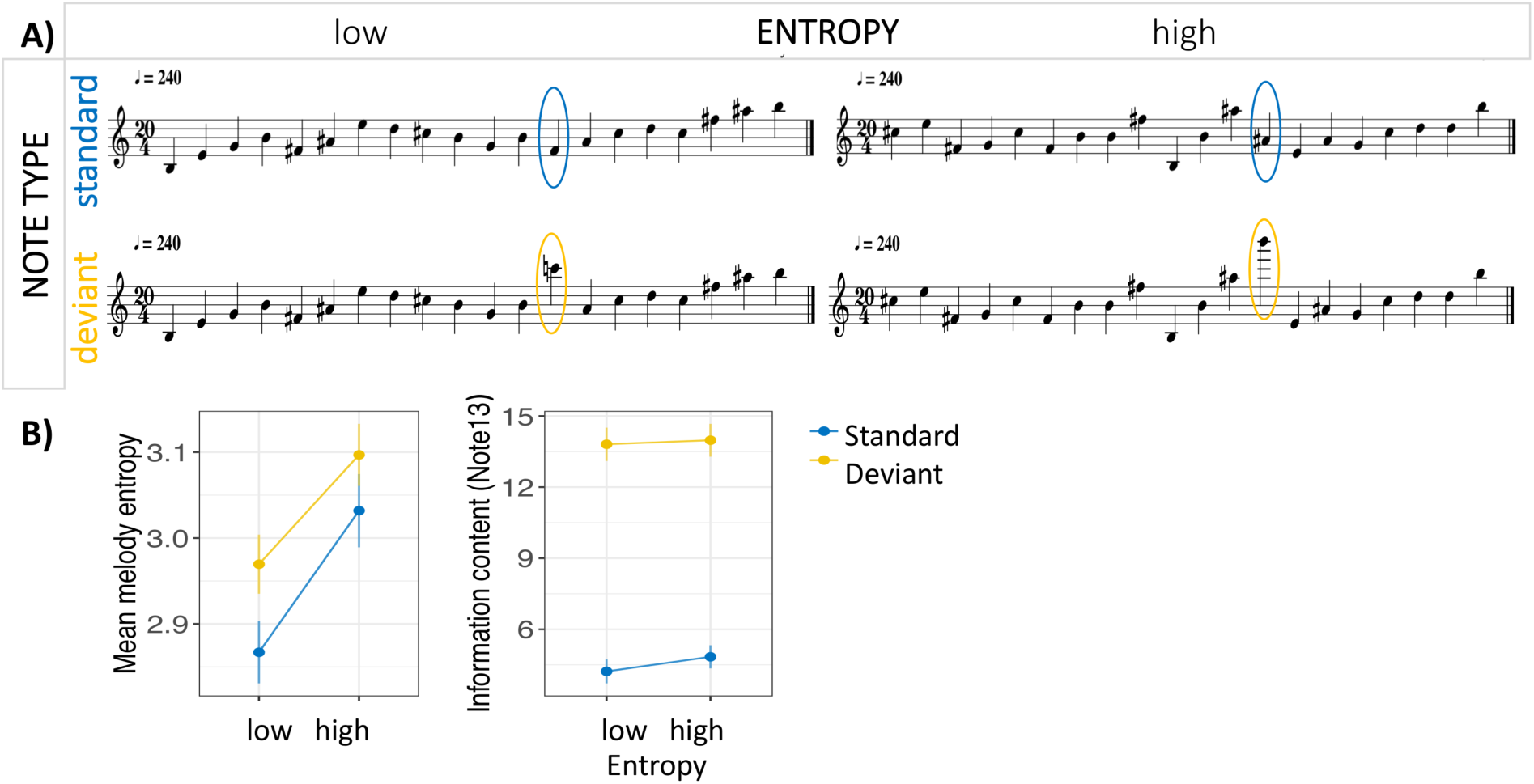
Experimental design. Participants listened to the melodies of 20 notes and had to rate how unexpected they found the last note on a seven-step Likert scale (– 1 equal ‘not at all unpredictable’ and 7 ‘extremely unpredictable’). The design included two factors: Contextual entropy (low/high) and Note type (standard/deviant). Melodies containing deviant notes where 50% of the trials and the deviant notes occurred always at the 13^th^ position. **B) Characterization of the stimuli**. A model of musical expectations was used to characterise the stimuli: Mean entropy was larger for high than low entropy melodies regardless of the presence of deviant (yellow) or standard (blue) tones. Mean information content of deviants (yellow) was larger than standard (blue) tones regardless of the entropy of the context.

## Methods

### Participants

Forty-seven participants (age: *M*=26.19, *SD*=6.24, min.=20, max.=52, 68% female, representing 15 nationalities) took part in the study. The sample scored relatively high on the general musical sophistication index, GoldMSI (Müllensiefen et al., 2014), with *M*=87.40, *SD*=25.24, min.=32, max.=120. A big sample size was chosen based on a previous experiment using musical stimuli (Laeng et al., 2016) and to ensure statistical power. As a post-hoc confirmation of the adequacy of our sample-size, we quantified effect sizes of the difference between PDR to deviants in high vs. low entropy contexts. The power analysis was conducted in the G*Power software package (version 3.1.9.2) with the setting p = .05 and N = 42 and confirmed that our sample size was adequate (1 - β > .9). Two participants reported ophthalmologic concerns or surgery prior to the experiment but were not excluded from participation as the pupil dilates even in blindsight participants (Weiskrantz, Cowey, & Barbur, 1999). Technical problems occurred during the recording of four participants, who were therefore excluded from the analysis. One subject was further excluded as blink gaps were too large to be interpolated. In sum, forty-two participants’ data were analysed.

Ethical approval for this study was granted by the Research Ethics Committee of Goldsmiths, University of London. Participants were instructed as to the purpose of the study, and consented to participate (written informed consent). Participation was remunerated with 5 pounds.

### Stimuli

One-hundred-twenty melodies were used in this study (thirty originally composed, thirty matched ‘shuffled’ versions, and sixty corresponding versions with a deviant tone always as the 13^th^ note). The melodic sequences comprised 20 tones, were 5 seconds long, isochronous (with an inter-onset-interval of 250 ms, 20/4 bar with 240 bpm), and had constant intensity and timbre (MIDI generated piano timbre).

The corpus of melodies was composed according to the principles of Western-tonal music and using all tones of the chromatic scale. Ambitus and tonal space varied across melodies. Interval size did not exceed a perfect fifth (Narmour, 2015) between adjacent notes. Thus, the original melodies were characterized by a smooth contour. To generate matched melodies that controlled for potential biases such as tonal space, pitch class, frequency range or ambitus, high entropy melodies were created from the original melodies by randomizing the order of constituent notes using MIDI processing tools (Eerola & Toiviainen, 2004). Our manipulation of entropy, whereby notes in original melodies were randomly shuffled without constraint, necessarily resulted in the mean absolute interval size being greater in high entropy than in low entropy melodies [L vs. H: t(58) = -2.644, p-value = .01]. Corpus studies of western tonal music show that large interval sizes are much less common than small ones (Huron, 2001). Thus, we anticipated that the presence of such large intervals would lead to the higher entropy levels desired for the high entropy condition.

Deviant notes were inserted in all 60 melodies (original and shuffled version) at the onset of the 13^th^ note (3000ms on the salient onbeat) in order to create the corresponding set of *deviant melodies* (Fig. 1). The deviant note was integrated into the second half of the melody in order to allow the establishment of an expectation-forming context before its occurrence. To ensure that differences in the PDR to the deviant notes between low and high entropy condition were not attributable to difference in the just preceding event, but to the context, the interval size between the deviant and the preceding 12th note was the same in matched pairs of high and low entropy melodies (see Fig. 1). However, interval size of the deviant varied between maj7 up/down, min9 up/down and aug11 up/down as those intervals sound particularly unusual within a melodic progression. Larger interval size of the deviant was assigned to matched pairs with lower entropy differences. This allowed for a variety of deviant interval sizes (a range of unexpectedness) while ensuring salience of deviants even in melodies showing relatively smaller entropy differences.

Entropy values were assigned by the IDyOM model to each note of the melodies. The IDyOM model considered one pitch viewpoint, namely ‘cpitch’, whereby chromatic notes count up and down from the middle pitch number (C=60) (Pearce, 2005). Through a process of unsupervised learning, the model was trained on a corpus (903 Western tonal melodies) comprising songs and ballads from Nova Scotia in Canada, German folk songs from the Essen Folk Song Collection and chorale melodies harmonised by J.S. Bach. The probability of each note of the stimulus used here were then estimated based on a combination of the training set’s statistics and those of the given melody at hand. The model outputs information content and entropy values. In mathematical terms, information content is inversely proportional to the probability of an event x_i_, with IC(x_i_) = -log2 p(x_i_) (MacKay, 2003), while the maximum entropy is reached when all potential events xi are equally probable, with p(x_i_) = 1/n, where n equals the number of stimuli. In psychological terms, information content represents how surprising each subsequent note is based on its fit to the prior context (Pearce & Wiggins, 2006). In contrast, entropy refers to the anticipatory difficulty in precisely predicting a subsequent note. Mean entropy values, obtained by taking the mean entropy of all notes, were used to characterise the predictability of each melody. Given our manipulation, mean entropy was largely, but not entirely, explained by the mean interval size of melodies (rho = .474, p < .001). These values were then used to predict subjective inferred uncertainty (measured as unexpectedness of last note) about each melody, and to categorise melodies into low and high entropy groups. By controlling that the interval preceding the deviant note was identical across high and low entropy condition, we predicted that a greater PDR to deviant notes in low than high entropy melodies should be attributable to the difference in the context.

### Procedure

The experiment was presented using the Experiment Builder Software and pupil diameter was recorded using EyeLink 1000 eye-tracker at a 250Hz sampling rate (SR Research, www.sr-research.com/experiment-builder). Prior to the data acquisition, a three-dot calibration was conducted to ensure adequate gaze measurements. Participants were further allowed to adjust the sound volume to a comfortable level and were asked to reduce head movements to a minimum throughout the recording session. As no differences were anticipated between the left and right pupil, the left pupil was recorded in ten and the right pupil in 32 participants depending on the participants’ dominant eye. To reduce motion artefacts, the head was stabilized using the SR Research Head Support chinrest placed 50 cm from the presentation screen.

During the experiment, the 120 melodies were presented binaurally through headphones in a randomized order. Each trial was triggered by the experimenter on the control computer when the fixation was stable at less than two arbitrary gaze units away from the fixation point. When recording was enabled, a white fixation dot on the grey screen turned black to prepare participants for the onset of the melody. The fixation cross was displayed for 7 seconds after stimulus onset. Each trial was preceded by a 400 ms baseline period and followed by a 2000 ms post stimulus offset. The melody was 5000 ms long and participants were instructed not to blink or move but to fixate during that whole period. Participants were allowed to take breaks to avoid fatigue effects at their convenience. A re-calibration procedure using the 3-dot-calibration was applied after each break.

Participants were instructed to carefully listen to the melody while fixating on the fixation point in the centre of the screen throughout the recording period. They were not informed about the deviant manipulation. After each trial, participants rated the final note on a Likert-scale ranging from 1 (not at all unpredictable) to 7 (extremely unpredictable) in a forced-choice task on the presentation screen. Data on the subjects’ musical expertise and sociodemographic background was collected at the end of each experiment using the GoldMSI (Müllensiefen et al., 2014). The whole study lasted approximately one hour.

### Data pre-processing

Blinks were identified and removed from the signal using MATLAB R2017b. These were characterized by a rapid decline towards zero from blink onset, and a rapid rise back to the regular value at blink offset. 100ms of the signal was removed before and after the missing data points (Troncoso, Macknik, & Martinez-Conde, 2008) and missing data were interpolated: four equally spaced time points were used to generate a cubic spline fit to the missing time points between blink onset (t2) and blink offset (t3) of the unsmoothed signal, with t1= t2-t3+t2 and t4= t3-t2+t3 (Mathôt, Fabius, Van Heusden, & Van der Stigchel, 2018). Trials containing more than 15% missing data were excluded from the analysis (*M*= .3, *SD* = .9 trials across subjects). Data were cleaned of artefacts using Hampel filtering (median filtered data; Pearson, Neuvo, Astola, & Gabbouj, 2016), and smoothened using a Savitzky-Golay filter of polynomial order 1 over the entire trial epoch. Finally, data were z-scored, and then baseline-corrected by subtracting the median pupil size of 400 ms baseline before melody onset. To analyse the time-window after deviant onset, data were baselined from 400 ms before deviant onset. The normalized pupil diameter was time-domain-averaged across trials of each condition.

### Experimental design and statistical analysis

We estimated a single time series for each sequence type: low or high entropy with either standard or deviant note type (*S-Low/D-Low/S-High/D-high*) by averaging across trials and participants. Statistical analysis was performed using Fieldtrip’s cluster-based permutation test (Maris & Oostenveld, 2007), with a significance threshold at 5% to control family-wise error-rate (FWER). This analysis revealed the time windows showing significant difference between the compared time series.

We first compared responses to deviant notes in the low and high entropy contexts with the corresponding standard notes using signals recorded across the entire melody duration (baselined 400 ms before melody onset). This ensured that differences between deviant and corresponding standards were not due to differences in the immediately preceding note. Then, to determine how the deviant PDR is affected by the entropy of the melodic context, we focussed on the time window from deviant onset to the end of the melody (3000 to 5000 ms epochs). We thus baselined to 400 ms before deviant onset, and we tested for an interaction of the deviant and context manipulation: (*D-Low – S-low*) vs. (*D-High – S-High*). Further, we compared the responses to standard tones in the two contexts (*S-Low – S-High*) to ensure that any differences were not driven simply by the standard notes (the control conditions).

To assess a potential influence of expertise on the PDR to deviants (data between 3000 and 5000 ms baselined to 400 ms before deviant onset), we computed the mean PDR to deviant trials as *D-Low – S- Low* for deviants in low entropy context, and *D-High – S-High* for deviants in high entropy context. Participants were split into two groups of musical experts and non-experts based on GoldMSI scores (*Mdn*=96). An ANOVA with within-subject factor context (low/high entropy) and expertise as between-subject factor (expert/non-expert) was computed.

## Results

### Stimuli characterization

Analyses were carried out to clarify the nature of all differences in information theoretic properties of the different stimuli. Figure 1B (left panel) shows the mean entropy values for all conditions (*D-Low*: *M* = 2.96, *SD* = .18; *D-High*: *M* = 3.09, *SD* = .19; *S-Low*: *M* = 2.86, *SD* = .19; *S-High*: *M* = 3.03, *SD* = .23). An ANOVA with context (low/high entropy) and note type (deviant / standard) as between-group factors yielded a main effect of context [*F*(1,116) = 15.20, *p* < .001, *np2* = .12], indicating higher entropy in *High* than *Low entropy* melodies [*t*(58) = 2.85, *p* = .006]. A significant main effect of note type [*F*(1,116) = 5, *p* = .027, *np2* = .04], was not supported by a further post hoc comparison [*t*(58) = -1.641, *p* = .11]. No interaction was found between the two factors [*F*(1,116) = .25, *p* = .617, *np2* < .01].

Figure 1B (right panel) shows that the unexpectedness of deviant notes, as reflected by information content values, was comparable between low and high entropy melodies. An ANOVA with between group factors context (low / high entropy) and note type (deviant / standard) yielded a main effect of note type [*F*(1,116) = 239.02, *p* < .001, *np2* = .67], a non-significant main effect of context [*F*(1,116) = .42, *p* = .517, *np2* = .01], and no interaction between the two factors [*F*(1,116) = .00, *p* = .714, *np2* < .01] – thus indicating higher information content in deviant than standard notes regardless of the context in which they were embedded [*t*(58) = -12.321, *p* < .001]. The similar IC levels of deviant for the two entropy conditions may be due to fact that in both low and high entropy contexts, deviants were similar in being characterised by very large interval departures away from the melodic contour (as opposed to the relatively naturalistic events that occurred in real melodies (Dean & Pearce, 2016). Critically, that deviant IC levels are similar for the two entropy conditions, supports our suggestion that stimulus context (and not just the IC level an incoming event) has the ability to modulate the PDR.

IC was positively predicted by interval size between the 12th and 13th note (*rho* = .862, *p* < .001), in line with research showing that amongst multiple psychological representations of pitch (e.g., height, contour, etc.), interval exerts a major contribution to perception of surprise (Levitin & Tirovolas, 2009; Pearce, 2018; Quiroga-martinez et al., 2019).

### PDR characterization

Figure 2A shows the time course of the PDR across conditions (*S-Low* = standard low entropy, *S-High* = standard high entropy, *D-Low* = deviant low entropy, *D-High* = deviant high entropy) baselined 400 ms before melody onset. A comparison between *S-Low* and *S-High* showed no difference in the PDR as a function of entropy levels of the melody, whilst the PDR to deviants (*D-Low* vs. *D-High*) was greater in predictable than unpredictable contexts (diverging at .56 s from deviant onset). The response to deviants in the predictable contexts was greater than the relative standard condition (*D-Low* vs. *S-Low*: *p* = .029), significantly diverging from *S-Low* between 3.57 and 5.64 s after melody onset. Conversely, the response to deviants in unpredictable contexts did not differ from the relative standard condition (*D-High* vs. *S-High)*, despite their high information content (see Figure 1B, middle panel).

**Fig. 2.**
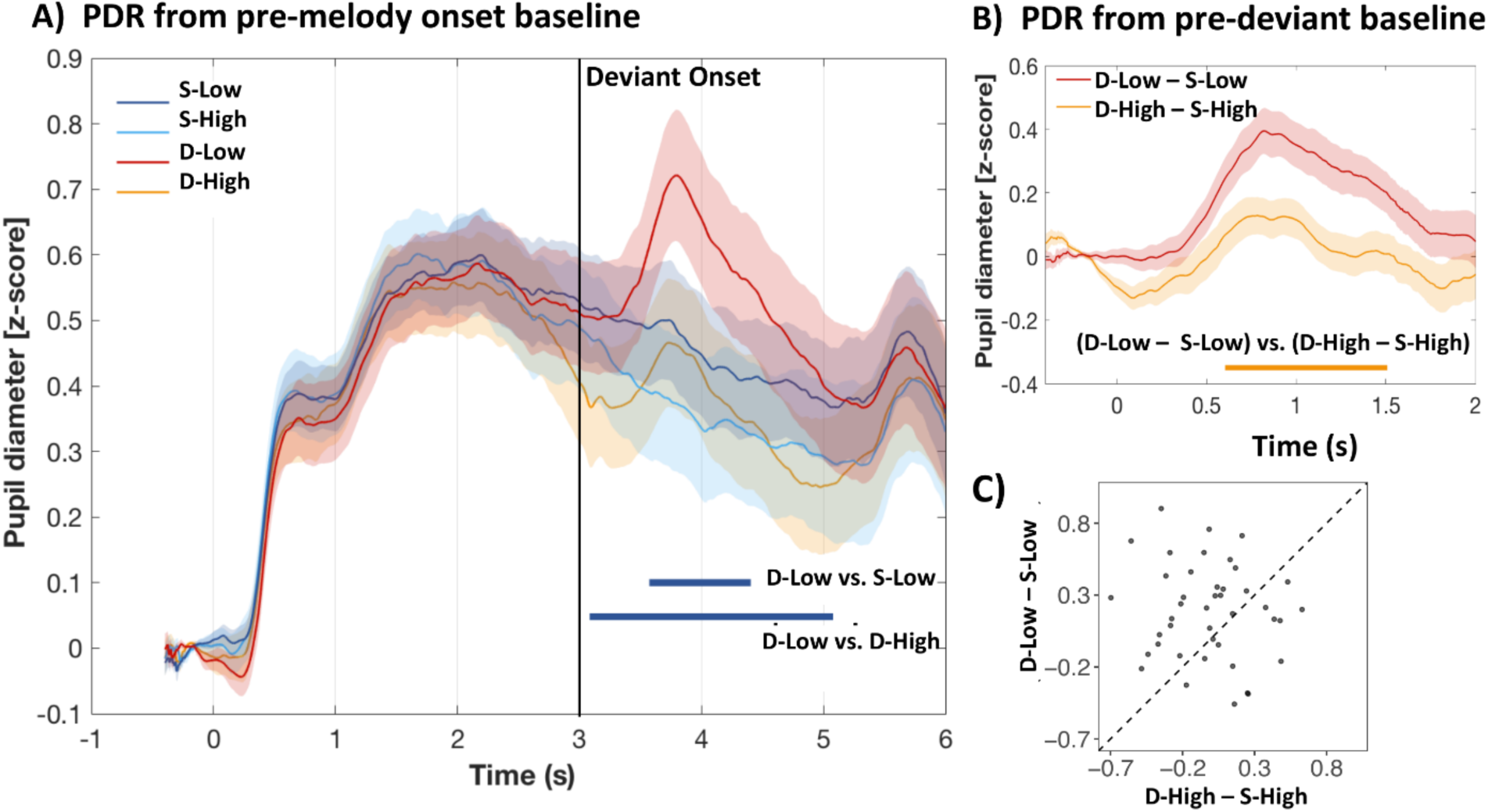
**A)** The PDR for all conditions from melody onset (*S-Low* = standard low entropy, *S-High* = standard high entropy, *D-Low* = deviant low entropy, *D-High* = deviant high entropy). The PDR to deviants compared to standard tones (*D-Low* vs. *S-Low*) in low entropy contexts diverged between 3.57 and 5.64 s from melody onset (.57 s from deviant onset), but did not differ in high entropy contexts. Also, the PDR to deviants was greater in low than in high entropy contexts (*D-Low* vs. *D-High*) (diverging at 3.056 s from melody onset), but there was no significant context-dependent difference between the standard tones (*S-Low* and *S-High*). Shaded regions around the curves represent standard error in the mean estimated with bootstrap resampling (1000 iterations; with replacement). **B)** Interaction effect of deviant and context entropy on the PDR after deviant onset. The difference between the PDR to deviant and standard tones was greater in low than in high entropy contexts. This effect emerged .59 after deviant onset and ended at 1.5 s. **C)** The relationship between the PDR for *D- High* – *S-High* and *D-Low* – *S-Low* conditions. Each data point represents an individual participant. Dots above the diagonal reveal that 67% of participants showed a greater PDR to deviants in low compared with high entropy contexts.

Figure 2B shows the PDR to deviants baselined 400 ms before deviant onset. We show the conditions *D-Low* and *D-High* following subtraction of the relative standard conditions. The comparison between the two time-courses confirmed that *D-Low* evoked a larger response than *D-High* starting at .59 s from deviant onset (*p* = .007) and ending at 1.5 s. Intrinsic noise in the baseline may explain the very early divergence between the curves, which however was not significant. Importantly, this pattern was observed in 67% of the participants (Figure 2C).

We run a post hoc analysis (Figure 3B) to investigate the relationship between the PDR related to the 13^th^ note (including both standard and deviant notes) and associated information content for low and high entropy melodies. For each subject and for each melody, the average PDR to the 13^th^ note was computed from the pre-deviant baseline (as in Figure 2B). A linear model with the factor context (Low /High) and information content as continuous variable was run to predict the PDR. This analysis yielded no main effect of context [*F*(1,116) = .005, *p* = .941, *np2* < .001] and a main effect of information content [*F*(1,116) = 4.911, *p* = .029, *np2* = .041] and a no significant interaction [*F*(1,116) = 3.274, *p* = .073, *np2* = .027]. This suggests that the PDR is sensitive to a large range of unexpectedness levels.

**Fig. 3.**
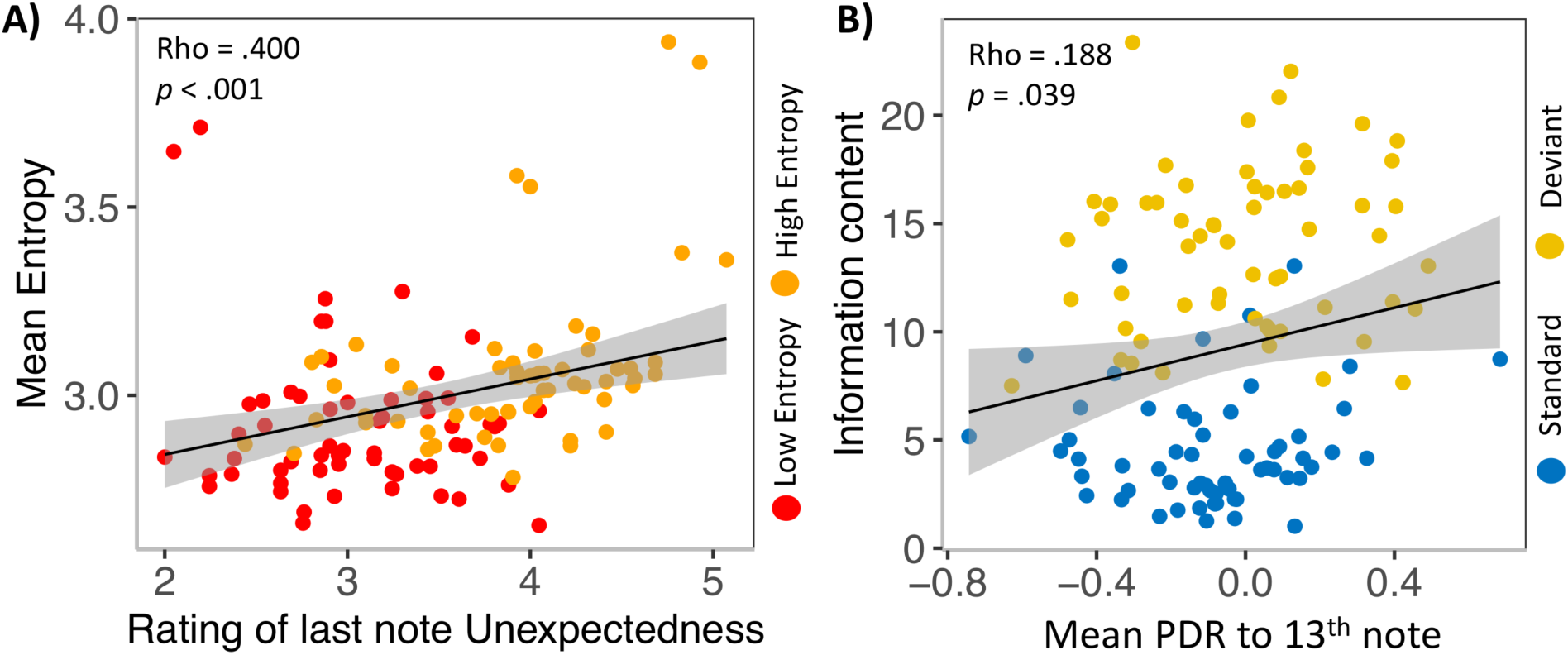
**A)** The stimulus entropy was computed by the IDyOM model, and validated by participants’ self-reports about the overall unexpectedness of each melody. Each dot represents one of the 120 melodies. Red and orange dots represent each of the low and high entropy melodies, respectively (*N* = 120). Shading represents s.e.m. **B)** The unexpectedness of the 13^th^ notes (deviants and standards) was estimated by the IDyOM model and showed a positive correlation (Spearman) with the mean evoked PDR. Blue and yellow dots represent each of the standard and deviant notes. Shading represents s.e.m.

We further investigated potential influences of expertise on the mean PDR to deviants (computed as *D-Low – S-Low* for deviants in low entropy contexts, and *D-High – S-High* for deviants in high entropy contexts on the data between 3000 and 5000 ms baselined to 400 ms before deviant onset as in Figure 2B). The ANOVA examining the effect of musical expertise on the mean PDR to deviants yielded a main effect of context [*F*(1,40) = 5.09, *p* = .03, *np2* = .11]. The main effect of expertise was not significant [*F*(1,40) = 3.72, *p* = .061, *np2* = .09], and no interaction between the two factors was seen [*F*(1,40) = 2.18, *p* = .147, *np2* = .05]. This confirmed that the mean PDR to deviants was greater in low than high entropy contexts although no considerable difference between experts and non-experts was observed.

Finally, we showed that the model reliably predicted subjective uncertainty levels (inferred entropy) of the melody progressions (Hansen & Pearce, 2014) (Figure 3A). The measure of the unexpectedness of the last note was collected after participants listened to each melody. We found that the mean ratings for each melody strongly correlated with the information content of the last note of the melody (*rho* = .355, *p* < .001), and with the mean IDyOM entropy values for that melody (*rho* = .400, *p* < .001). This analysis validated our categorization of melodies based on the IDyOM output.

## Discussion

We report that pupil dilation response (PDR) to behaviourally irrelevant deviants occurs when deviants are embedded in predictable rather than unpredictable melodies. We showed that the amplitude of the response is predicted by the information content (or unexpectedness) of the musical deviants. We also replicate the previous finding that listeners’ experience of uncertainty is predicted by the entropy of the music (Hansen & Pearce, 2014). These results show that the same sudden environmental change leads to differing levels of arousal depending on whether it occurs in low or high states of uncertainty. Our results suggest that the more stable predictions formed in predictable rather than unpredictable contexts may be more abruptly violated by surprising events, possibly leading to greater changes in the listeners’ arousal state.

The observed modulatory effect of context predictability on the PDR to deviants is consistent with a body of electrophysiological work showing context effects on mismatch like responses at the cortical level (Garrido et al., 2013; Quiroga-Martinez et al., 2019; Southwell & Chait, 2018). Here we show a similar pattern but in autonomic markers of arousal, as reflected by the PDR. Pupil response is thought to be driven by norepinephrine activity in the locus coeruleus (LC) (Joshi, Li, Kalwani, & Gold, 2016). This is a key subcortical nucleus which widely connects to the brain (Sara, 2009) to signal unexpected and abrupt contextual changes (Alamia et al., 2019; Damsma & van Rijn, 2017; Zhao et al., 2019). It has been hypothesised (Zhao et al., 2019) that MMN-related brain systems (Garrido et al., 2009; Hsu et al., 2015; Southwell & Chait, 2018) may trigger norepinephrine-mediated updating or interruption of ongoing top-down expectations. In line with this hypothesis, Alamia et al. (2019) have shown a correlation between pupillary response and MMN-like response evoked by violations of expectations, providing evidence of a link between the sources of these two responses. Top-down expectations about unfolding sensory signals have been associated with the temporo-frontal network in music-violation paradigms, which is thought to link present and past information to generate predictions about forthcoming events (Bianco, Novembre, Keller, Seung-Goo, et al., 2016; Koelsch, Gunter, et al., 2002; Tillmann, Janata, & Bharucha, 2003). Based on this existing evidence and in line with a model resetting hypothesis, our results suggest that listeners generate stronger predictive models (in the predictable melodies). These may be supported by temporo-frontal cortical regions, and require a greater signal (greater PDR) to be interrupted.

Albeit indirectly, our results also establish a link between subcortical and cortical activity in response to unexpected events under different states of uncertainty. Increased MMN response under low states of uncertainty (Quiroga-Martinez et al., 2019; Southwell & Chait, 2018) are replicated in the pupil response, reflecting a general increase in arousal. A possible interpretation, in line with the predictive coding theory (Friston, 2005), is that more stable expectations, representative of a strong predictive model, are reflected in precision-weighing of the prediction error, and hence a stronger response when the input mismatches the current predictions. Conversely, in high entropic contexts predictive models are weak, and the prediction error attenuated. This mirroring pattern between cortical (MMN) and subcortical (as reflected by the PDR) responses may have important behavioural advantages. Specifically, strong predictive models can suddenly be abandoned when they are revealed to be erroneous, thus allowing speedy reorienting behaviours and quicker engagement with new potentially relevant stimuli. One limitation with regard to this possible interpretation is the nature of the deviant events used here, whereby the deviating event was a single note that did not lead to any long-lasting changes in the statistics of the unfolding sequence. Further studies combining cortical and pupil response measurements in continuously changing stimuli are necessary to corroborate our working hypothesis.

The absence of difference between high and low entropic contexts in the sustained pupil response (in the first half of the melodies) suggests that the pupil is relatively unresponsive to slowly unfolding stimulus structures, at least when listeners are not required to actively track them (Alamia et al., 2019; Zhao et al., 2019). Whilst cortical responses have been shown to be sensitive to the statistics of the unfolding stimulus structure (Barascud, Pearce, Griffiths, Friston, & Chait, 2016; Sohoglu & Chait, 2016; Southwell et al., 2017), subcortical responses may be less sensitive. This suggests they may be more vulnerable to stimulus properties and tasks demands (Zhao et al., 2019).

We also found that the PDR mismatch response is positively predicted by unexpectedness of incoming notes (in line with electrophysiological studies (Omigie et al., 2013; Quiroga-martinez et al., 2019), but it seems not to be modulated by degree of musical expertise. Larger MMN responses have been shown for musicians in a range of studies examining electrophysiological correlates of expectancy violation in music (Koelsch, Schmidt, & Kansok, 2002b; Oechslin, Van De Ville, Lazeyras, Hauert, & James, 2013; Tervaniemi, Tupala, & Brattico, 2012). One possibility is that this reflects a ceiling effect whereby the rather salient deviants used here were relatively easy to detect (given their large interval departures away from the melodic contour). Future studies examining the PDR to more subtle differences in musical structure may be expected to show similar expertise effects to those reported in previous studies.

Finally, our results provide evidence of music’s usefulness in investigating the neural mechanisms underlying processing of stimuli statistical properties in a common, highly structured, and ecologically valid type of auditory stimulus, as music. While our work here focused on pitch expectations, previous studies have shown that music-induced temporal expectations are also tracked by the PDR (Damsma & van Rijn, 2017). Future experiments could address, for example, how introducing rhythm to concurrent melodic lines may affect the PDR to unexpected events. Similarly, that music induced chills – high arousal physiological responses associated with subjective pleasure – are associated with increased PDR (Laeng et al., 2016) suggest a usefulness of music in examining the relationship between stimulus information theoretic properties and reward processing. Whilst the IDyOM model used here is only able to deal with monophonic MIDI music, the development of the model for polyphonic music is underway (https://psyarxiv.com/wgjyv/) and it is expected that the approach we take here will be beneficial in a wider range of contexts in future years. By showing that predictive uncertainty can be used to modulate prediction-error related arousal, our findings have implications for understanding the variety of forms listeners’ aesthetic appreciation of music may take. However, considering, more generally, the tight coupling between the error-related norepinephrine system and the reward-seeking dopaminergic pathway (Laeng et al., 2016; Xing, Li, & Gao, 2016; Zatorre & Salimpoor, 2013), our results emphasize that measuring the PDR may be useful for investigating the reward value of information across a range of modalities and domains.

In sum, we show that pupillometry in the auditory domain can reliably track the effect of context uncertainty on responses to sudden environmental change and independently from overt behavioural responses. Given the tight interplay between cortical and subcortical mechanisms involved in precision weighted anticipatory processing, a first milestone is set towards the non-invasive quantification of related arousal responses.

## Author Contributions

**RB** conceived the experiments; analysed the bulk of the data; wrote the manuscript.

**EP** conceived and performed the experiments; contributed to analysis; wrote the manuscript; secured the funding.

**DO** conceived the experiments; contributed to analysis; wrote the manuscript; secured the funding.

## Acknowledgments

We are grateful to Prof. Maria Chait and Sijia Zhao for inspiring discussions.

## Data Availability Statement

The datasets for this study can be found in the OSF repository (link: https://osf.io/u2qdr/?view_only=7a9fa6bacbb249b090282377c1542d29).

## Informed Consent Statement

All participants provided written informed consent. The study was approved by the Ethics Committee at Goldsmiths, University of London.

